# Profiling of chromatin accessibility across *Aspergillus* species and identification of transcription factor binding sites in the *Aspergillus* genome using filamentous fungi ATAC-seq

**DOI:** 10.1101/857284

**Authors:** Lianggang Huang, Xuejie Li, Liangbo Dong, Bin Wang, Li Pan

## Abstract

To identify *cis*-regulatory elements (CREs) and motifs of TF binding is an important step in understanding the regulatory functions of TF binding and gene expression. The lack of experimentally determined and computationally inferred data means that the genome-wide CREs and TF binding sites (TFBs) in filamentous fungi remain unknown. ATAC-seq is a technique that provides a high-resolution measurement of chromatin accessibility to Tn5 transposase integration. In filamentous fungi, the existence of cell walls and the difficulty in purifying nuclei have prevented the routine application of this technique. Herein, we modified the ATAC-seq protocol in filamentous fungi to identify and map open chromatin and TF-binding sites on a genome-scale. We applied the assay for ATAC-seq among different *Aspergillus* species, during different culture conditions, and among TF-deficient strains to delineate open chromatin regions and TFBs across each genome. The syntenic orthologues regions and differential changes regions of chromatin accessibility were responsible for functional conservative regulatory elements and differential gene expression in the *Aspergillus* genome respectively. Importantly, 17 and 15 novel transcription factor binding motifs that were enriched in the genomic footprints identified from ATAC-seq data of *A. niger*, were verified in vivo by our artificial synthetic minimal promoter system, respectively. Furthermore, we first confirmed the strand-specific patterns of Tn5 transposase around the binding sites of known TFs by comparing ATAC-seq data of TF-deficient strains with the data from a wild-type strain.

## Introduction

There have been 339 species identified in the *Aspergilli* genus (Samson et al. 2014), and they are of broad interest because of their industrial applications, importance as pathogens for animals and crops, and relevance for basic research. *Aspergillus* section Nigri is the fungal genus with the most sequenced genomes, as genomes from 27 species are already publicly available (Vesth et al. 2018), and genomic analyses have led to a better understanding of fungal biology and improvements in the industrial use of these organisms (Pel et al. 2007; Andersen et al. 2011; Yin et al. 2014). Gene regulation is of major way by which fungi physiologically act in environmental circumstances and respond to changing conditions (Todd et al. 2014; Schape et al. 2019). However, in comparison with *Saccharomyces cerevisiae*, the knowledge of genome-wide regulatory mechanisms in *Aspergilli* is lagging. Transcription factors (TFs) are key regulators of biological processes that function by binding to transcriptional regulatory regions to control the expression of target genes. The CREs related to TF binding play pivotal roles in the regulation of gene expression, and each TF recognizes a collection of similar DNA sequences, known as binding site motifs (Felsenfeld 1992; Qiu et al. 2016). Therefore, the ability to identify these CREs and TF motifs throughout *Aspergillus* genomes is an important step in understanding the regulatory functions of TF binding and gene expression. Analysis of genome sequences has revealed an average of 600 putative transcription factors in each *Aspergillus* fungus genome (Galagan et al. 2005; Pel et al. 2007). Nevertheless, approximately 5% of the TFs in the *Aspergillus* genus have only been identified and have not been studied further (Galagan et al. 2005; Pel et al. 2007). Most genome-wide TF binding sites in the *Aspergillus* genome have not been experimentally determined or computationally inferred, and the sites remain unknown.

The gold-standard method for identifying *in vivo* CREs for TFs of interest is chromatin immunoprecipitation DNA sequencing (ChIP-seq) (Johnson et al. 2007). However, the scarcity of sequence-specific antibodies for most *Aspergillus* TFs or tagged target proteins for the more than 600 TFs in *Aspergillus* has prevented the widespread application of this method in *Aspergillus* (Park 2009). So far, only a few ChIP-seq experiments have used epitope-tagged or customized TF antibodies to study *Aspergillus* TFs, including CrzA (de Castro et al. 2014), SrbA (Chung et al. 2014), and MetR (Amich et al. 2013) of *Aspergillus fumigatus*; CRE-1 of *Neurospora crassa* (Sun and Glass 2011); Tri6 of *Fusarium graminearum* (Nasmith et al. 2011); and *MoCRZ1* of *Magnaporthe oryzae* (Kim et al. 2010). This finding reveals that the available filamentous fungi ChIP-seq data are not abundant; further, performing ChIP-seq experiments in filamentous fungi is full of challenges. Additional high-throughput sequencing methods, i.e., DNase-seq and formaldehyde-assisted isolation of regulatory elements with sequencing (FAIRE-seq) combine enzymatic digestion of isolated chromatin with sequencing, were developed to identify CREs on a genome-wide scale (Giresi et al. 2007; Wang et al. 2015). The shortcomings of these methods are that they require millions of cells as starting material, complex and time-consuming sample preparations, and many potentially loss-prone steps, such as adaptor ligation, gel purification and cross-link reversal (Zentner and Henikoff 2014). For these reasons, the DNase-seq analysis that we previously carried out in *Aspergillus oryzae* could not be effectively replicated (Wang et al. 2015). Therefore, the development of feasible and scalable methods is required to facilitate the identification of regulatory elements.

Assay for transposase accessible chromatin sequencing (ATAC-seq) is a recently developed technique used to identify accessible regions and DNA footprints (Buenrostro et al. 2015; Li et al. 2019). Tn5 transposase integrates sequencing adapters directly into DNA, eliminating the need for multiple reactions and purification steps typically required for construction of a sequencing library. As a result, significantly lower amounts of starting nuclei are required for the investigation of CREs. ATAC-seq is now routinely being applied to systematically identify *cis-*regulatory regions and DNA footprints in humans (Buenrostro et al. 2013), mice (Wu et al. 2016), zebrafish (Fernandez-Minan et al. 2016), *Drosophila* (Bozek et al. 2019), *Caenorhabditis elegans* (Daugherty et al. 2017), *Arabidopsis thaliana* (Tannenbaum et al. 2018), *Saccharomyces cerevisiae* (Schep et al. 2015) and so on. However, in *Aspergillus* species, the existence of cell walls and the purification of nuclei preparation prevented the routine application of this technique. To overcome this obstacle, we performed an ATAC-seq assay for *Aspergilli* by protoplast preparation under cultivation conditions. We combined ATAC-seq and RNA-sequencing of different *Aspergillus* species and TF mutants to profile chromatin accessibility and identify gene regulatory elements *in vivo* on a genome-wide scale. Furthermore, we characterized the identified CREs *in vivo* by the synthetic minimal promoter driving expression of the reporter gene. Collectively, this study is an important step towards the meticulous analysis of industrial microbial transcriptional regulatory networks and provides valuable resources for future research aimed at characterizing or using gene regulatory elements.

## Results

### The ATAC-seq assay in filamentous fungus

To apply the ATAC-seq assay to identify chromatin accessibility and regulatory elements in filamentous fungi, we developed an ATAC-seq protocol for *Aspergillus species* by adding a step that released native nuclei by detergent lysis of protoplasts prior to the transposition step (Fig. 1A). A total of 5×10^4^ protoplasts were harvested and subjected to the extraction of native nuclei in lysis buffer containing 0.05% IGEPAL CA-630 (Sigma, I8896). After Tn5 transposition, PCR and electrophoresis were performed as previously described (Buenrostro et al. 2013); then, discrete nucleic acid bands clearly appeared in the gel image (Fig. 1B), indicating Tn5 tagmentation of regions between nucleosomes. The insert size distribution of sequenced fragments from *A. niger* chromatin had a clear periodicity of approximately 200 bp, suggesting many fragments are protected by integer multiples of nucleosomes (Fig. 1C).

**Figure 1.**
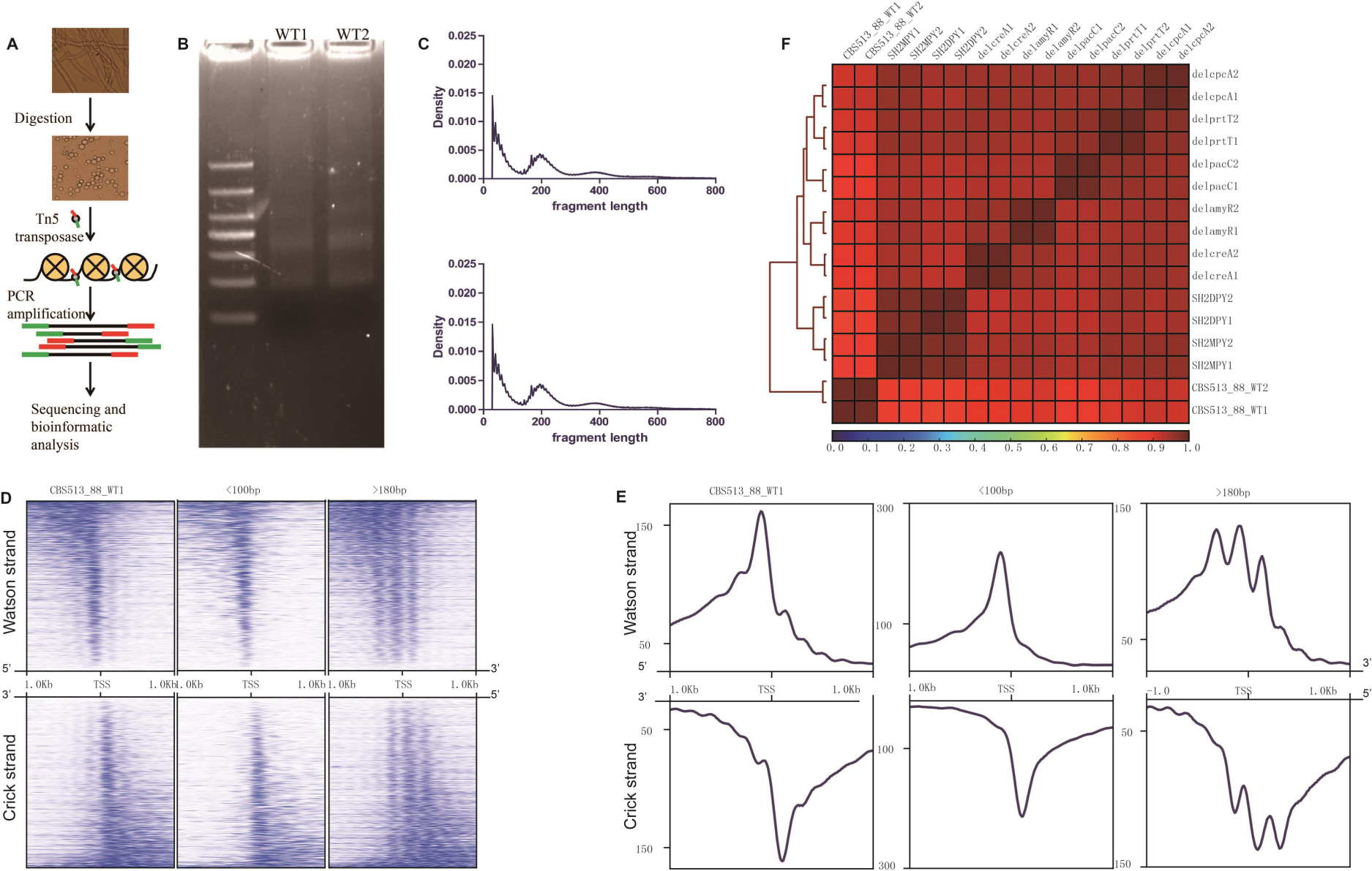
Schematic illustration of the Assay for Transposase-Accessible Chromatin with high-throughput sequencing (ATAC-seq) in *Aspergillus niger*. (A) Protoplasts used for nuclei isolation and schematic procedure of ATAC-seq technique. Protoplasts were prepared under the cultivation condition by adding lywallzyme. After filtration via miracloth membrane, the concentration of extracted protoplasts reached 10^5^/μL. (B) Fragment sizes for amplified ATAC-seq libraries by 15 cycles of PCR reaction. 10^5^ protoplasts of *A. niger* CBS513.88 wildtype strains (WT1 and WT2) were used for Tn5 transposase reaction, respectively. (C) Insert sizes determined by high-throughput sequencing. Peaks with different fragment length indicate that the fragments contain one or more nucleosomes. (D) Heatmaps showing the distribution of accessible regions around TSSs in *A. niger* CBS513.88. (E) Read coverage of the 1-kb region flanking TSSs in *A. niger* CBS513.88. (F) Heatmap clustering of Spearman correlation coefficients for 16 *A. niger* ATAC-seq datasets.

We analysed the pattern of mapped ATAC-seq reads around the TSSs in *A. niger* CBS513.88 WT1. Heatmaps showed the intensity of mapped ATAC-seq reads flanking 1-kb upstream and downstream of annotated TSSs in the AspGD database (Fig.1D). Furthermore, separate heatmaps showed that fragments less than 100 bp clustered immediately upstream of TSSs throughout the *A. niger* genome. Fragments between 180 and 247 bp, corresponding to nucleosome footprints, were depleted from TSSs throughout the genome. Overall, the identified accessible regions were mostly enriched around the TSSs (Fig. 1D), which is consistent with the fact that these regions usually contain active regulatory *cis-*elements (Qiu et al. 2016). The signal intensity of accessible regions became weaker as the distance from the TSS became greater in both the Watson and Crick strands (Fig. 1E), indicating that different regulatory factors might combine with *cis*-elements in the Watson and Crick strands to control the expression of the corresponding genes. Furthermore, using ATAC-seq for *A. niger*, we generated paired-end ATAC-seq libraries for 16 samples, and the libraries ranged from 35.9 to 104.9 million reads (Supplemental Table S3). The percentages of reads in two replicate samples of CBS513.88_WT1 and CBS513.88_WT2 that aligned to the *A. niger* CBS513.88 genome were 84.23% and 86.31%, respectively (Supplemental Table S3). The insert size distributions of sequenced fragments from 16 ATAC-seq libraries of *A. niger* had a similar periodicity (Supplemental Fig. S3). Spearman correlation analysis was used to inspect the reproducibility of biological and technical replicates. Heatmap clustering of Spearman correlation coefficients for 16 *A. niger* ATAC-seq datasets revealed a strong correlation between replicates of the same strain or the mutant and a lower correlation between different strains (Fig. 1F). Taken together, these data suggested that the ATAC-seq protocol of filamentous fungus *A. niger* could provide data on accessible regions of chromatin in a genome-wide search for regulatory elements.

### Identification of genome-wide chromatin accessibility and comparison of open chromatin profiles among *Aspergillus* species and strains

Syntenic analysis showed that most of the genomic sequences were homologous between *A. niger* strains CBS513.88 and SH2, except for a missing DNA fragment in *A. niger* CBS513.88 contig VIII_An18 and a fragment in *A. niger* SH2 contig S6 (Fig. 2A). Visualization of the *A. niger* CBS513.88 and SH2 ATAC-seq data sets in a control eye on a genome-scale revealed that they were similar to each other, and the Tn5 integration regions of naked genomic DNA were uniformly distributed along the entire chromosome (Fig. 2A). For *the A. niger* SH2 ATAC-seq data, there was indeed no ATAC-seq signal in the missing DNA fragment of *A. niger* CBS513.88 contig VIII_An18 (Fig. 2A). A total of 7297 and 7697 accessible chromatin regions (peaks) were identified in *A. niger* CBS513.88 and SH2, respectively (Fig. 2B, Supplemental Table S4). Among these accessible regions, 6524 regions overlapped, indicating that these regions contained common *cis-*elements for both strains. These common regulatory elements might play a basal transcriptional regulatory role in *A. niger* species. Additionally, 773 and 1173 accessible regions are specific for *A. niger* CBS513.88 and SH2, respectively, and these accessible regions contained differential *cis-*elements for each strain. A heatmap of the peak signal for the 2-kb flanking region of the peak centre (Fig. 2C) showed that most of the accessible regions formed sharp peaks in the centre and ranged from narrow to wide, which provided valuable data depending on *cis*-element motifs in these accessible regions. On the genome scale, accessible regions are highly enriched in promoter regions (Fig. 2D), which was consistent with the distribution of accessible regions in Fig. 1D. In addition to promoter regions, the TTS regions contain the maximally accessible regions (12.88% for CBS513.88 and 15.70% for SH2, Fig. 2D). These accessible regions in TTSs indicate that gene downstream regions also contain regulatory elements for gene transcription.

**Figure 2.**
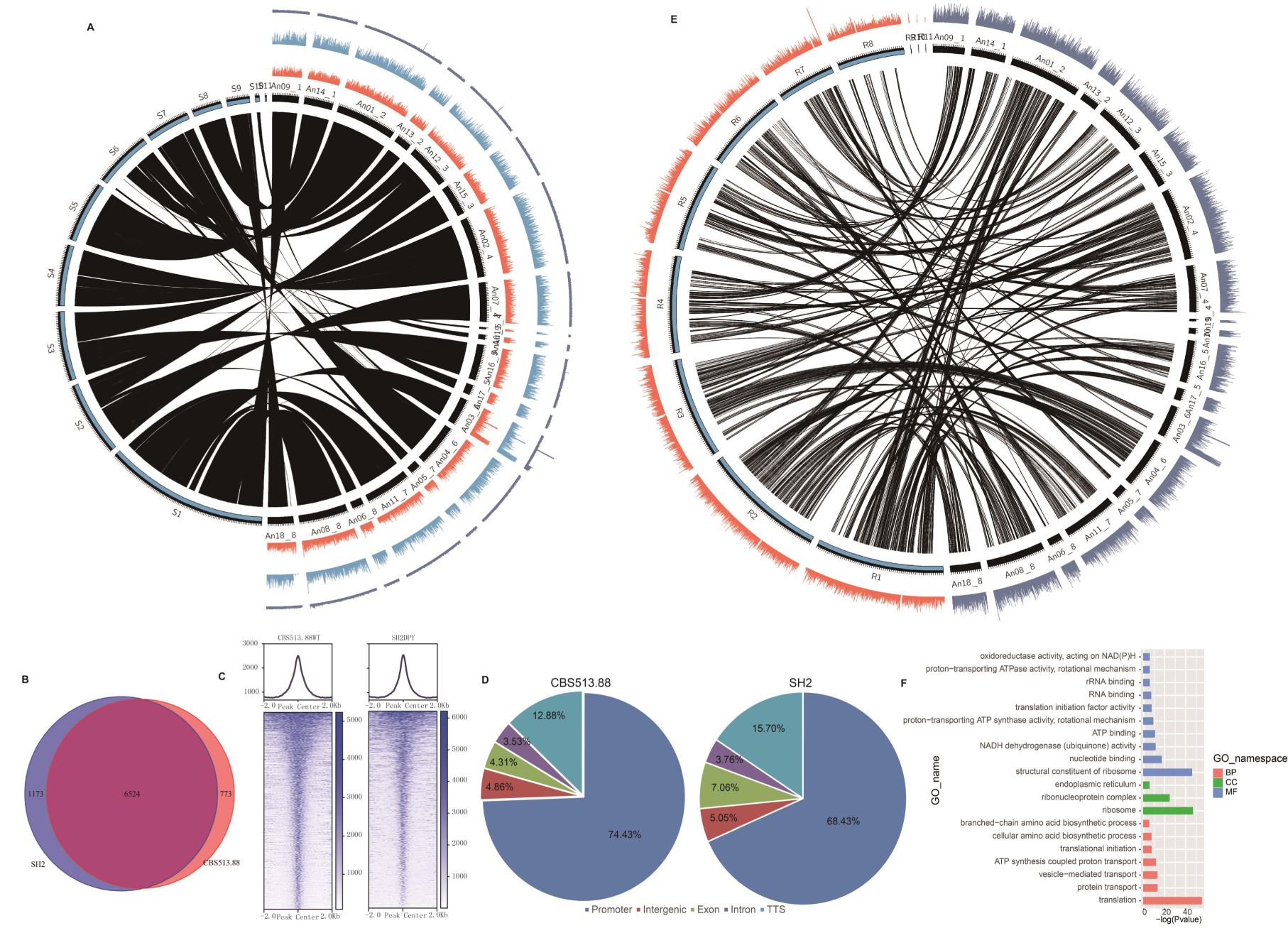
Whole-genome landscape of chromatin accessibility in *A. niger*. (A) Syntenic analysis between *A. niger* CBS513.88 (contig An01-An19) and SH2 (contig S1-S11), and visualization of ATAC-seq peak signals in *A. niger* CBS513.88 (orange) and SH2 (blue). *A. niger* CBS513.88 naked genomic DNA was used as the integration control of Tn5 transposase. (B) A Venn diagram showing the overlap of accessible regions between *A. niger* CBS513.88 and SH2. (C) Density plots of mapped reads (upper) and heatmap of peak length (bottom) for the 2-kb region flanking the peak center. (D) Distribution of accessible regions in the promoter, intergenic region, exon, intron, and TTS in *A. niger* CBS513.88 and SH2. (E) Syntenic analysis of ATAC-seq peaks between *A. niger* CBS513.88 (contig An01-An19) and *A. oryzae* niaD30 (contig R1-R8). (F) Functional clustering of overlapping ATAC-seq peaks of *A. niger* CBS513.88 and *A. oryzae* niaD300.

To understand the syntenic orthologues of accessible regions across the *Aspergillus* species, we compared the *A. niger* CBS513.88 and *A. oryzae* niaD300 ATAC-seq peaks on a genome-wide scale. Our results showed that these two *Aspergillus* species had strong peak synteny, and there were 1564 syntenic orthologues of ATAC-seq peaks identified between *A. niger* CBS513.88 and *A. oryzae* niaD300 (Fig. 2E, Supplemental Table S5). Then, we located the target genes of these syntenic peaks and analysed their GO enrichment function (Fig. 2F). Ribosomes related to GO categories were significantly enriched in these target genes, especially for the following three categories: “structural constituent of ribosome (MF)”, “ribosome (CC)” and “translation (BP)”. These highly enriched target genes indicated that *Aspergillus* species strains possessed a powerful capacity in protein synthesis and energy-based protein transportation, which make them suitable for use as microbial cell factories. These data suggest that our ATAC-seq protocol for a filamentous fungus could identify chromatin accessibility in regulatory elements on a genomic scale.

### Identification of significantly changed chromatin accessibility driven by transcription factors and chromosome regulatory factor in *Aspergillus* species

A clustering map of ATAC-seq peaks for all 16 *A. niger* samples indicates that there were common ATAC-seq patterns among all *A. niger* strains and their TF-defective strains (Fig. 3A). However, there were obvious differences between the *A. niger* CBS513.88 and SH2 strains. *A. niger* SH2 and its derivatives did not have ATAC-seq signals corresponding to the 200-kb missing genomic DNA fragments (Fig. 3A) (Yin et al. 2014). To investigate the changes in chromatin accessibility between *A. niger* SH2 and its TF-defective strains, we identified the ATAC-seq peaks that significantly changed accessibility (FDR<0.05) among *A. niger* SH2 cultured under different conditions and its TF-defective strains (Fig. 3B and Supplemental Table S6). A total of 4009 of the differential ATAC-seq peaks in the promoter regions were identified when comparing *A. niger* SH2 cultured in DPY and MPY media (Supplemental Table S6). We found that changing the culture conditions could slightly alter the differential chromatin accessibility, in which 3853 of the differential ATAC-seq peaks were specific to *A. niger* SH2 cultured under MPY medium (Supplemental Table S6). Almost all upregulated genes were driven by the MPY-specific ATAC-seq peaks (Fig. 3B: MPY_vs_DPY). Furthermore, we identified TF-specific chromatin accessibility using the peak signal intensity of *A. niger* SH2 compared with the peaks observed in TF-defective strains (FDR < 0.05). In total, 1641 of the amyR-specific peaks, 1156 of the prtT-specific peaks, 773 of the cpcA-specific peaks, 1301 of the pacC-specific peaks, and 1305 of the creA-specific peaks in the promoter regions exhibited significantly higher accessibility (FDR<0.05) than those in the corresponding TF-defective strains (Supplemental Table S6). The distributions of upregulated genes were unbalanced and trended towards TF-specific chromatin accessibility (Fig. 3B).

**Figure 3.**
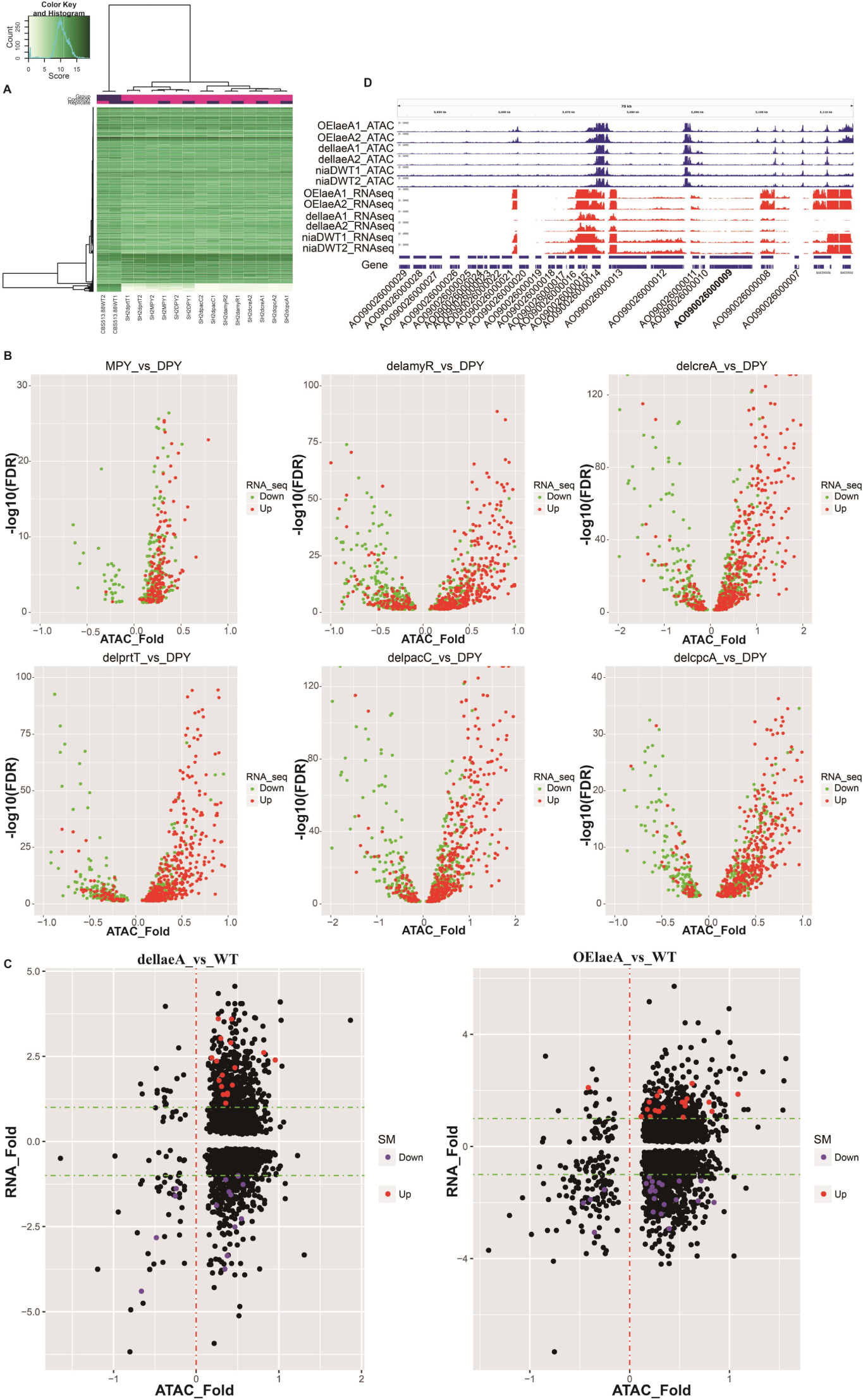
Differential open chromatin accessibility analysis of *Asperigllus*. (A) Clustering analysis of ATAC-seq peaks for 16 *A. niger* samples. (B) Volcano plot of differentially expressed genes (|log2(Fold_Change)|≥1, P<0.05) corresponding to differential chromatin accessibility regions (FDR<0.05) among *A. niger* strain and TF-deficient strains. ATAC_Fold>0 means an increase in chromatin accessibility. (C) Scatter plot of differentially expressed genes (P<0.05) and chromatin accessibility between *A. oryzae* WT strain and laeA mutants. Significantly differentially expressed SM genes (|log2(Fold_Change)|≥1, P<0.05) are indicated by red and purple dots. The green dotted line indicates |log2(Fold_Change)|=1. (D) A representative aflatoxin biosynthetic pathway in *A. oryzae* showing a comparison between ATAC-seq and RNA-seq for samples of *A. oryzae* WT strain and laeA mutants.

The biosynthetic gene clusters of secondary metabolites (SMs) in *Aspergillus* species are frequently silent or inactive under normal laboratory conditions due to a tendency to be located in subtelomeric regions with a high degree of heterochromatin condition; however, these genes can be activated by changing the chromosomal state of the SM gene clusters from heterochromatin to euchromatin (Brakhage 2013). LaeA is a key regulatory factor involved in histone methylation that mediates the process of reversing the heterochromatin status, thereby changing the chromatin landscape of a large genomic region (Bok and Keller 2004; Bok et al. 2006). To investigate whether the changes in chromatin accessibility activated SM biosynthetic gene clusters, we constructed *A. oryzae* niaD300 dellaeA and OElaeA mutants and analysed the corresponding libraries with ATAC-seq (Supplemental Table S3, Supplemental Fig. S4) and RNA-seq. We identified the ATAC-seq peaks with significantly changed accessibility (FDR<0.05) between the dellaeA mutant or the OElaeA mutant and the *A. oryzae* niaD300 wild-type strain (Fig. 3C and Supplemental Table S7). There were 4725 and 5572 differential ATAC-seq peaks identified from *A. oryzae* OElaeA and dellaeA mutants compared with the *A. oryzae* niaD300 wild-type strain, respectively. SM biosynthetic gene clusters in the *A. oryzae* RIB40 genome contain 621 biosynthetic SM genes predicted by Secondary Metabolite Unique Regions Finder (SMURF) (Khaldi et al. 2010). The chromatin regions in front of 138 and 164 of the SM biosynthetic genes in *A. oryzae* OElaeA and dellaeA mutants significantly changed accessibility (FDR<0.05) to induce significant regulation of 42 and 32 SM biosynthetic genes, respectively (Fig. 3C and Supplemental Table S7). The 54 backbone genes of SM biosynthesis include polyketide synthases (PKSs) and non-ribosomal peptide synthetases (NRPSs) from the *A. oryzae* RIB40 genome; only 12 backbone genes of SM biosynthesis significantly changed accessibility (FDR<0.05) to induce significant regulation of the corresponding SM backbone genes in OElaeA and dellaeA mutants, which contained 7 NRPSs, 4 PKSs, and 1 DMAT (Supplemental Table S7). We found that the knockout and overexpression of the *laeA* gene only up- and downregulated the SM biosynthetic genes that were expressed in the wild-type strain, while there was never *de novo* activation of those SM biosynthetic genes that were not expressed in the wild-type strain (Supplemental Fig. S5). Furthermore, we investigated chromatin accessibility (ATAC-seq peaks) and transcriptional activity in the famous aflatoxin (AF) biosynthesis pathway of *A. oryzae* (Fig. 3D). The sequences of the aflatoxin (AF) biosynthesis gene clusters were divided into two parts: active and inactive chromatin regions. The sequences from AO090026000004 to AO090026000020 were the active chromatin regions, which were characterized by highly expressed biosynthesis genes. The over-expression and knockout of the *laeA* gene were able to change the chromatin accessibility in the active regions. The key transcription factor aflR (AO090026000014) was in the region of high chromatin accessibility, while the coactivator aflJ (AO090026000015) became highly expressed following the over-expression of *laeA*. The other sequences from AO090026000020 to AO090026000031 were in inactive chromatin regions in which no ATAC-seq peaks were identified, and biosynthesis genes had no expression or the lowest level of expression. The aflatoxin (AF) biosynthesis pathway components were located in a laeA-dependent SM biosynthetic gene cluster. Taken together, the reason for the lack of AF biosynthesis in *A. oryzae* might be inactive chromatin regions around the AF biosynthesis pathway.

### Identification of footprints in filamentous fungus *Aspergillus niger*

In addition to identifying open chromatin regions, the ATAC-seq assay can be applied to reveal DNA footprints, which can be used to identify potential TF-binding sites *in vivo* (Lu et al. 2017; Karabacak Calviello et al. 2019; Li et al. 2019). To systematically identify genome-wide DNA footprints, we used the “*wellington_footprints.py*” function of the pyDNase package (Piper et al. 2013), and we identified Tn5 integration-insensitive sites from samples of *A. niger* CBS513.88 and SH2 strains and footprints from ATAC-seq peaks (candidate accessible regions) identified by MACS2 (Fig. 4A). Only the footprints within ATAC-seq peaks were identified as ATAC-seq footprints. In total, there were 24,925 and 22,548 DNA footprints detected *in vivo* across the *A. niger* genome from 7,297 and 7,697 ATAC-seq peaks from *A. niger* CBS513.88 and SH2 ATAC-seq data, respectively (footprint data from all wild-type and TF-defective strains can be found in Supplemental Table S8). A total of 14,176 DNA footprints overlapped between *A. niger* CBS513.88 and SH2 ATAC-seq data. The DNA footprints identified by the ATAC-seq data were enriched 1-kb upstream and 0.5-kb downstream of the TSSs (Fig. 4B). To further characterize genomic footprints, we performed *de novo* motif enrichment analysis on the footprints from *A. niger* CBS513.88 and SH2 using the “*findMotifsGenome.pl*” function of the HOMER package (Heinz et al. 2010) using default settings. A total of 15 and 17 *de novo* TF binding motifs with a P-value threshold of 1.0E-12 were enriched more than 5-fold from the two sets of footprint data from *A. niger* CBS513.88 and SH2, respectively (Fig. 4C and Supplemental Table S9). *De novo* TF binding motifs of other SH2 TF-defective strains with a P-value threshold of 1E-50 were also identified using the HOMER method (Supplemental Fig. S6 and Table S9). The E-box of bHLH factors (motif TCACGTGATC), the SltA homologue binding motif (motif CTGCCTGAGGCA), and the core element of bZIP factors (motif GCTGAGTCAGCV) were among the top 5 most enriched motifs according to P-value in all *de novo* TF binding motifs (Fig. 4C). Instances of each TF binding motif were identified using the “*annotatePeaks.pl* -m” function of the HOMER package to search for DNA sequences in the 24,925 DNA footprints from *A. niger* CBS513.88 and the 22,548 DNA footprints from *A. niger* SH2 (Supplemental Table S10). Instances of genes controlled by *de novo* TF binding at motifs were identified in footprints in the promoter regions, and they were analysed for the functional enrichment of corresponding TF binding motifs (Supplemental Fig. S7). In the *A. niger* SH2 strain, 269 genes under the control of the *de novo* motif TCACGTGATC (E-box) were enriched in GO categories “structural constituent of the ribosome” and “translation” (Supplemental Fig. S7A). In addition, 154 genes under the control of the *de novo* motif CTGCCTGAGGCA (SltA homologue) were enriched in the molecular function GO categories “ATP binding” and “structural constituent of the ribosome”; in the cellular components of “ribosome”, “mitochondrial inner membrane”, and “vacuole”; and for the biological processes of “translation” and “proteolysis” (Supplemental Fig. S7B). Then, 54 genes under the control of the *de novo* motif GCTGAGTCAGCV (bZIP factors) were also enriched in the molecular function GO categories “zinc ion binding”, “DNA binding”, and “transcription factor activity” in the nucleus for the process of regulation of transcription (Supplemental Fig. S7C).

**Figure 4.**
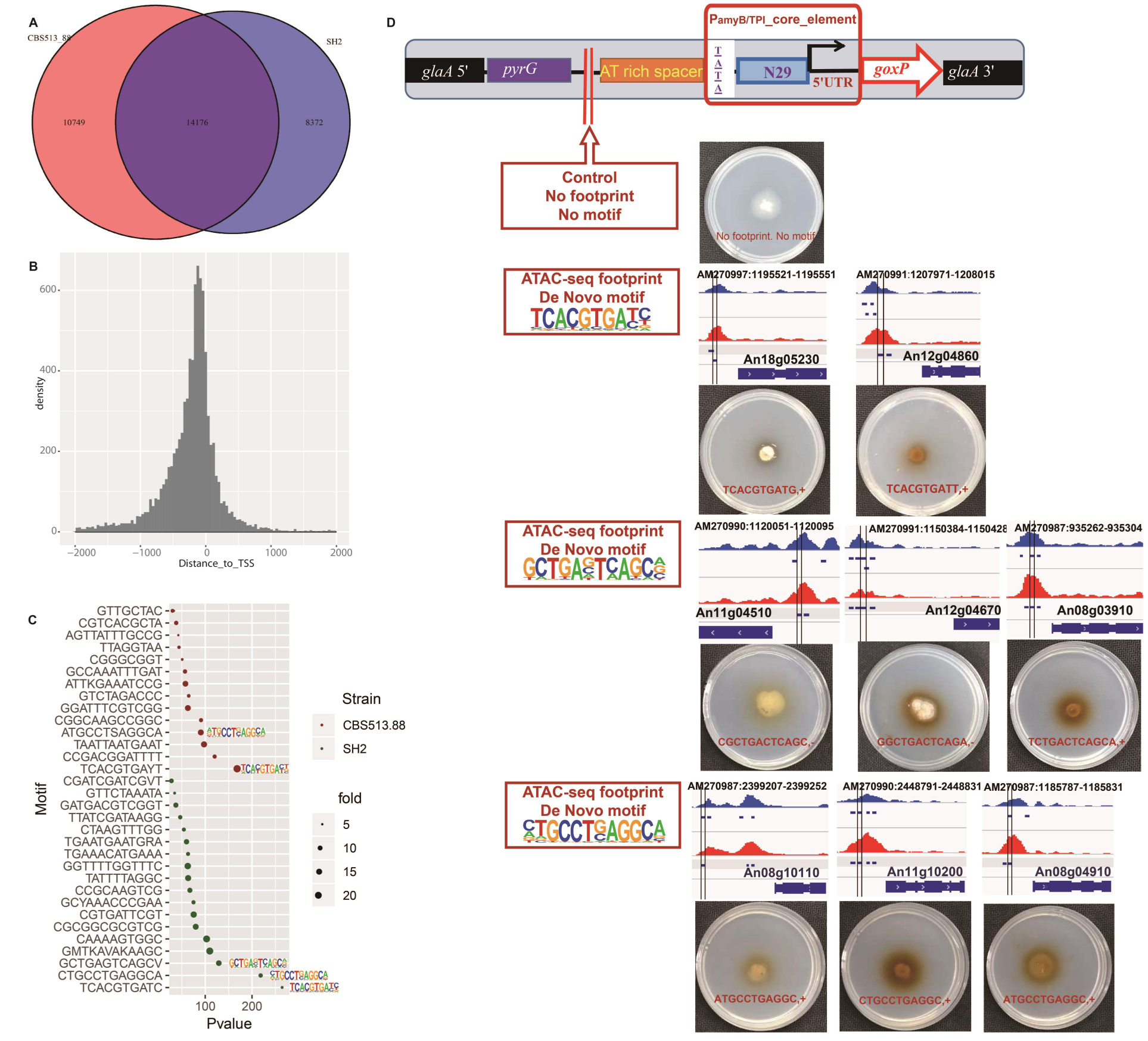
Identification of *de novo* predicted footprints in *A. niger* CBS513.88 and SH2, and functional verification in regulating gene expression. (A) Venn diagram analysis of identified footprints in *A. niger* CBS513.88 and SH2. (B) Statistical analysis of the distances between the identified DNA footprints and the TSSs. (C) *De novo* prediction of transcription factors in *A. niger* CBS513.88 (red) and SH2 (green). The circle size represents the enrichment fold. The top three transcription factor motifs in *A. niger* SH2 are labeled with the corresponding binding motifs. The x-axis represents P-value. (D) *In vivo* functional verification of putative *de novo* transcription factor targeting sites were carried out by artificially synthesized minimal promoters. The red dividing line represents the location of the footprints to be verified. From top to bottom, there were 2 E-box of bHLH factors (motif TCACGTGATC), 3 core elements of bZIP factors and 3 SltA homologs binding motif (motif CTGCCTGAGGCA) target sites validated by phenotype plates *in vivo*.

Furthermore, to detect whether the ATAC-seq footprints containing the TF binding motifs drove the expression of functional target genes, we constructed a minimal synthetic promoter consisting of a core element, an ATAT-binding box, an AT-rich spacer, and a footprint library-driven UAS element with a TF binding site motif (Supplemental Fig. S2). A 29-bp sequence from the TSS in the amyB (An05g02100) promoter of *A. niger* was chosen as the core element due to the high-level expression of the *amyB* gene. The glucose oxidase *goxC* (An01g14740) gene was used as a reporter, the expression of which would be revealed by a coloured colony. The repression under ER stress (RESS) element identified in our prior study (Zhou et al. 2015) was used as a positive control, and it also produces coloured colonies (Supplemental Fig. S2). As a negative control, a sequence containing only the *amyB* core element, the ATAT sequence, and the AT-rich spacer was constructed (no coloured colonies but basal transcription can occur (Supplemental Fig. S2)). We used this minimal promoter system to determine whether the ATAC-seq footprints containing *de novo* motifs, such as motif TCACGTGATC (E-box), motif CTGCCTGAGGCA (SltA homologue) and motif GCTGAGTCAGCV (bZIP factors), could drive the expression of target genes (Fig. 4D). Verifying the translation and ribosome functions controlled by the motif TCACGTGATC (E-box), the footprint of An12g04860, which encodes a ribosomal protein of the large subunit L30, was found to drive the expression of the reporter gene *in vivo,* resulting in coloured colonies (Fig. 4D). The footprints containing motif CTGCCTGAGGCA located in the promoter regions of An08g04910 and An11g10200, which encode the NADH-ubiquinone oxidoreductase and cytochrome C oxidase subunit in the electron respiratory chain, respectively, also drove the expression of the reporter gene *in vivo*. The footprints containing the motif GCTGAGTCAGCV, which is located in the promoter regions of An11g04510 and An12g04670 (related to translation release factor activity and translation initiation factor, respectively), were confirmed to drive the expression of the reporter gene *in vivo*. Taken together, genome-wide DNA footprints were systematically identified from accessible chromatin regions based on our ATAC-seq data. In addition, *de novo* motifs enriched from ATAC-seq footprints were able to drive the expression of their target genes *in vivo*.

### Known TF motifs of *Aspergillus niger* enrichment in ATAC-seq footprints

We determined the transposase Tn5 integration patterns around the binding sites of known TFs such as AmyR, PrtT, PacC, and CreA in the *A. niger* SH2 strain cultured under MPY medium; we also determine the patterns in naked genomic DNA subjected to ATAC-seq to control for Tn5 transposase sequence bias *in vitro* (Fig. 5A). While we observed Tn5 integration patterns in the motif sequence of binding sites from four known TFs in our ATAC-seq data, these patterns for known TFs were also evident in naked genomic DNA. However, we found that distributions of Tn5 integration sites in the flanking regions of binding site motif sequences from known TFs in our ATAC-seq data from the *A. niger* SH2 wild-type strain (MPY medium) were asymmetrical when comparing the anti-sense and sense strand; further, the patterns which were completely different from the symmetrical patterns observed in the naked genomic DNA. Furthermore, to elucidate whether strand-specific Tn5 integration patterns around TF binding sites are caused by protein/DNA interactions, we constructed TF-defective strains of *amyR*, *prtT*, *pacC*, and *creA* and determined the distribution of Tn5 integration sites around four known TF binding sites, which controlled for Tn5 transposase sequence bias *in vivo* (Fig. 5A). In this experiment, there was no corresponding known TF to be expressed *in vivo*, so TF binding sites should not exist. Tn5 integration sites around the known TF binding sites in the absence of the corresponding TF condition showed a similar pattern to that of genomic DNA, which was a symmetrical integration pattern around TF binding sites on both DNA strands. The asymmetrical pattern of Tn5 integration around TF binding sites in our ATAC-seq data was distinctly different from the no TF-binding pattern on both TF-defective strains *in vivo* and naked genomic DNA *in vitro* (Fig. 5A). We demonstrated here that strand-specific Tn5 integration patterns around TF binding sites are caused by protein/DNA interactions. This strand-specific asymmetrical signal in the flanking regions of TF binding sites could be used as a signal indicating an active TF binding site.

**Figure 5.**
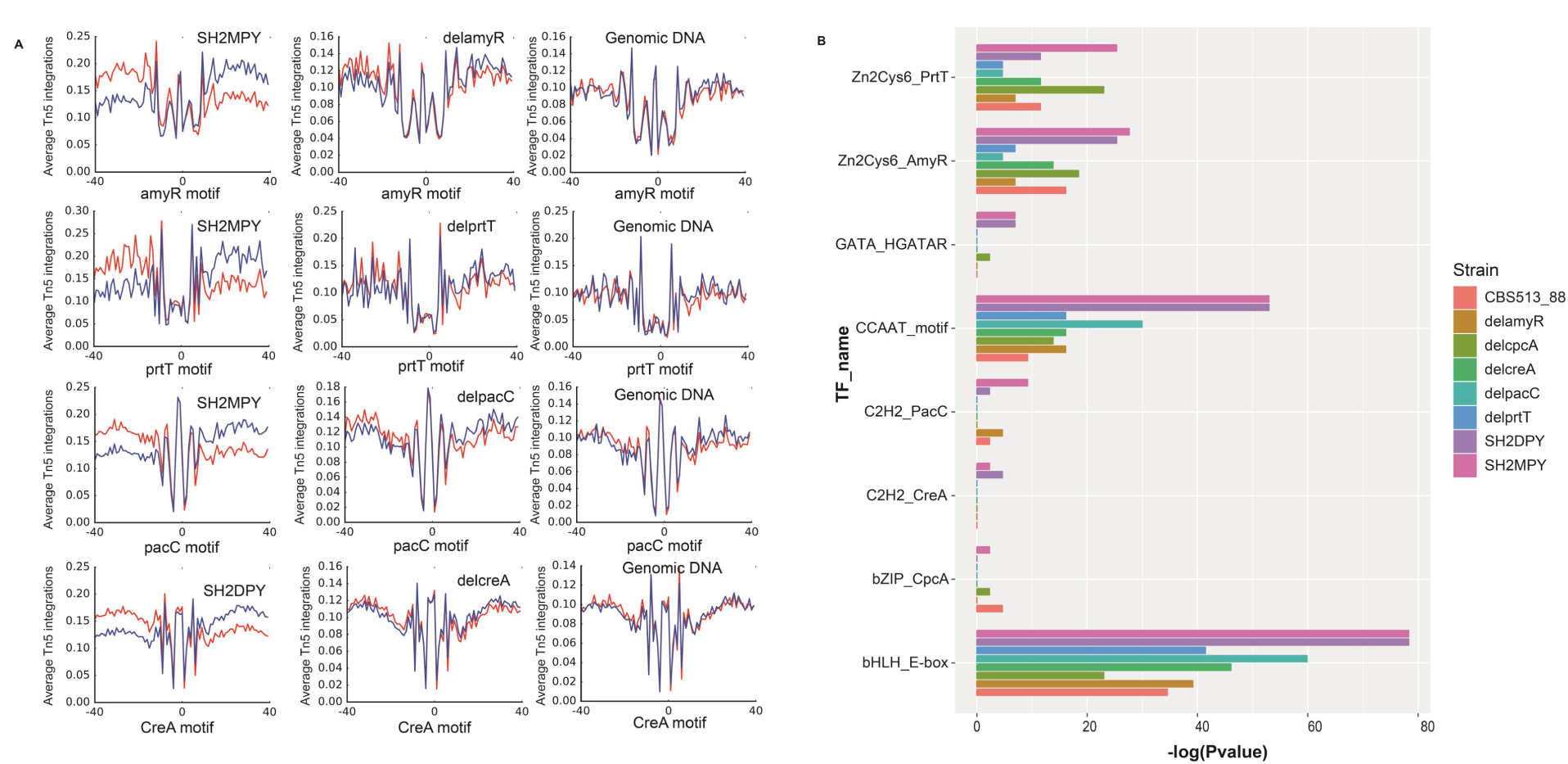
Distribution of Tn5 integration around known footprints and enrichment analysis of known TF motifs. (A) Distribution of Tn5 integration sites in the ATAC-seq data of SH2MPY (left), TF deletion strains (middle) and naked genomic DNA (right) around known motifs including *amyR*, *prtT*, *pacC,* and *creA*. Tn5 integration sites showed similar patterns around known TF motifs between TF deletion strains and naked genomic DNA whereas the patterns are different for SH2MPY. The red line indicates the integration sites on the positive-sense strand whereas the blue line indicates those on the anti-sense strand. (B) Enrichment analysis of known TF motifs in *A. niger* CBS513.88, SH2 and TFs deletion mutants; -log(P-value) was used as a significant criterion.

To determine the enrichment of known TF binding motifs from ATAC-seq footprints by comparing ATAC-seq data from *Aspergillus niger* wild-type strain and its TF-defective strains, we collected the available binding motif sequences of 19 known *Aspergillus* TFs (Wang et al. 2015) and employed the “findMotifsGenome.pl” function of the HOMER package, which we used to identify overrepresented motifs of known *Aspergillus* TFs from ATAC-seq footprint data. We found that the binding motifs of bZIP CpcA, C2H2 CreA, and PacC were not overrepresented in the ATAC-seq footprints of the corresponding TF-defective strains (P-value >1.0E-2) (Fig. 5B). Although the binding motifs of the Zn_2_Cys_6_ family AmyR and PrtT were also enriched in the footprints from the TF-defective strains delamyR and delprtT, respectively, the enrichment P-values of AmyR and PrtT motifs were less than 1.0E-2 (Fig. 5B). The binding motifs for bHLH family E-box, CCAAT binding complex (CBC), Zn_2_Cys_6_ family AmyR, and PrtT were significantly enriched in footprints from ATAC-seq data across all *A. niger* strains, including CBS513.88, SH2, and TF-defective strains; the enrichment was especially clear in the *A. niger* SH2 strain that was cultured in MPY medium (Fig. 5B). Other TF family motifs, such as bZIP CpcA, GATA AreA, C2H2 CreA, and PacC, were only enriched in the ATAC-seq footprints of the *A. niger* strain SH2 cultured in MPY and DPY medium (Fig. 5B). Footprints that contained a significant motif match were considered to be instances of predicted binding motifs. The TSS that was nearest to the motif of each known TF was defined as the target gene for that TF footprint or TF binding motif. We used the “annotatePeaks.pl -m” function of the HOMER package to detect the motif instances of known TFs from footprints by comparing *Aspergillus niger* wild-type strains and the corresponding TF-defective strains (Supplemental Table S11). The motifs of AmyR, PrtT, PacC, and CreA were identified in 765, 158, 821, and 2865 motif instances in the *Aspergillus niger* SH2 strain cultured in MPY medium, respectively (Supplemental Table S11). However, we still detected 593 instances of AmyR binding, 103 instances of PrtT binding, 656 instances of PacC binding, and 2648 instances of CreA binding from footprints in the TF-defective strains delamyR, delprtT, delpacC, and delcreA, respectively (Supplemental Table S11).

Furthermore, to investigate whether instances of TF-binding drove the expression of target genes, we used TF-binding motif sites in ATAC-seq footprints combined with RNA-seq data for the changes in gene expression between the *Aspergillus niger* SH2 strain cultured in MPY medium and the corresponding TF-defective strains (Fig. 6A). The expression of 1248 genes in the amyR-deficient strain (delamyR) and 1242 genes in the prtT-deficient strain (delprtT) was downregulated compared to what was seen in the control of the *A. niger* SH2 strain (Fig. 6A). Only 78 and 23 genes downregulated in the amyR-deficient and prtT-deficient strains overlapped with motif target genes AmyR and PrtT, respectively (Fig. 6A). As expected, 23 target genes under the control of PrtT were mainly involved in serine-type peptidase activity, and 78 target genes under the control of AmyR were functionally enriched in hydrolase activity, acting on glycosyl bonds (Fig. 6B). A total of 1644 genes were upregulated in the creA-deficient strain (delcreA) compared with the control of the *A. niger* SH2 strain, in which 269 genes overlapped with motif target genes from creA (Fig. 6A). GO functional enrichment analysis revealed that 269 genes under the control of CreA were primarily involved in the molecular functions of hydrolase activity and transporter activity; further, the enriched genes were structural constituents of the ribosome and were cellular components of membranes and ribosomes, and their biological processes involved carbohydrate catabolism, carbohydrate transport, and translation (Fig. 6B).

**Figure 6.**
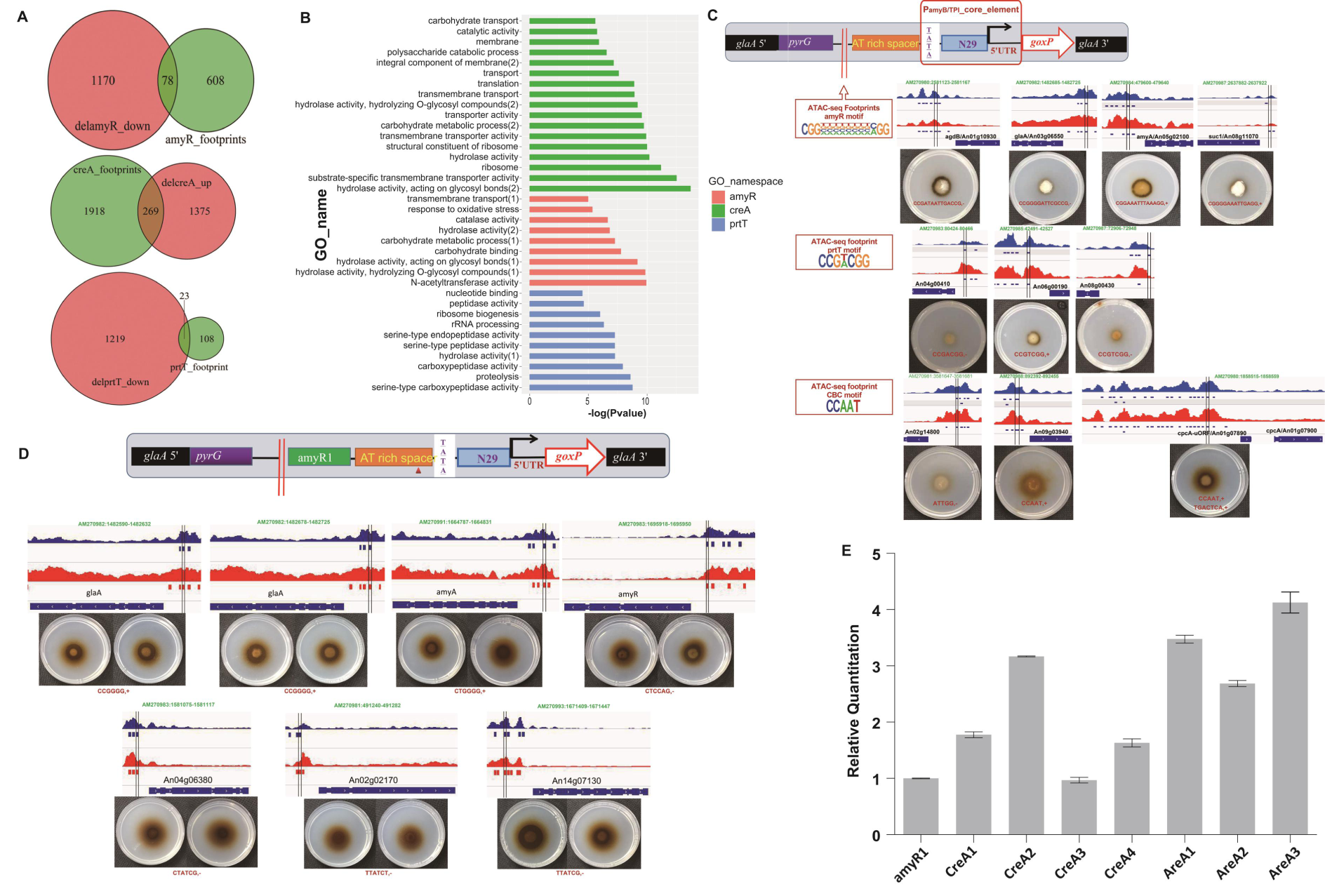
Functional analysis and verification of known TF motifs in ATAC-seq footprints. (A) Venn diagram analysis of TF binding motif instances of ATAC-seq footprints combined with RNA-seq data; (B) Functional classification of the intersection genes of TF binding motif instances of ATAC-seq footprints combined with RNA-seq data through FungiFun. (C) *In vivo* functional verification of putative known transcription factor targeting sites was carried out by artificially synthesized minimal promoters. The red dividing line represents the location of the footprints to be verified. From top to bottom, there were 4 AmyR, 3 PrtT and 3 CBC box (CCAAT) target sites validated by phenotype plates *in vivo*. (D) The repressors, CreA and AreA, were functionally verified by an artificial promoter containing AmyR instances. The active sites of 4 CreA and 3 AreA were verified by phenotype plates, respectively. (E) qRT-PCR analysis of the *goxC* gene driven by CreA and AreA targeted instances.

To experimentally assess whether footprints with these known TF binding motifs that were identified by ATAC-seq data were functional TF binding sites, we performed footprint-driven reporter assays *in vivo* (Fig. 6C). Three footprints contained the AmyR binding motif, and they were located in promoters of glycosyl bond hydrolase genes: An01g10930 (*agdB*), An03g06550 (*glaA*) and An05g02100 (*amyA*). These AmyR binding motifs drove the expression of the reporter gene, resulting in coloured colonies. Three footprints containing the PrtT binding motif in the promoter of peptidase genes, including An04g00410 (dipeptidyl peptidase III), An06g00190 (Sedolisin family secreted protease) and An08g00430 (serine-type carboxypeptidase), also drove the expression of the reporter gene. Furthermore, the footprints containing the short CCAAT binding motif in the promoters of the target genes An02g14800 (*pdiA*) and An09g03940 were able to drive the expression of the reporter gene to form coloured colonies.

In addition to identifying the activation function of ATAC-seq footprints containing the known TF binding motifs, we further investigated whether ATAC-seq footprints containing repressor binding motifs could downregulate the expression of target genes (Fig. 6D). We constructed a minimal synthetic promoter with the footprint-driven UAS elements containing the AmyR binding motif instance (An01g10930, *agdB*) as a positive control (Fig. 6D). The footprint containing the repressor binding motif was assessed by assembling downstream of the AmyR footprint. *A. niger* CreA (An02g03830, contains two Cys2His2 zinc finger motifs) is the transcriptional regulator that mediates carbon catabolite repression (Drysdale et al. 1993). However, our results showed that footprints containing CreA binding motif sites located in promoters of glycosyl bond hydrolases such as *glaA* (An03g06550), *amyA* (An12g06930), and *amyR* (An04g06910) were able to upregulate the expression of the reporter gene (Fig. 6D). Two footprints containing the CreA binding motif in the *glaA* promoter possessed the same ability to activate functional gene expression (Fig. 6D). We also assessed the other repressor, AreA, that controls nitrogen utilization. The same results showed that footprints containing the AreA binding motif could also drive the upregulation of the reporter gene (Fig. 6D). Furthermore, RT-PCR results also confirmed that footprints with repressor CreA and AreA binding motifs upregulated reporter gene expression levels (Fig. 6E). Taken together, the transcriptional regulators CreA and AreA act not only as repressors but also as activators.

## Discussion

In ATAC-seq, Tn5 transposase integrates its adaptor payload into locations of open chromatin, which can be amplified by PCR and assessed with high-throughput sequencing to study protein: DNA interactions on a genome scale (Buenrostro et al. 2013). Here, we developed an ATAC-seq protocol for filamentous fungi to identify chromatin accessibility on a genome scale. Our initial attempts failed to generate ATAC-seq data from the chromatin following digestion of Tn5 transposase, harvesting the mycelium and reducing it to powder by grinding it in liquid nitrogen with a mortar and pestle (Gonzalez and Scazzocchio 1997). The gel image analysis shows no discrete nucleic acid bands, and this method resulted in a high noise-to-signal ratio after sequencing; these results are most likely due to the presence of a chitinous cell wall, which limits the accessibility of external nucleases to nuclei. In our modified *Aspergillus* ATAC-seq protocol, the lysis of the *Aspergilli* cell wall was carried out by adding hydrolase during cultivation, and protoplasts were obtained by filtration through a miracloth membrane. High-quality intact nuclei released from *A. niger* protoplasts, which are similar to mammalian cells, were used to improve the data collected from ATAC-seq. ATAC-seq requires few cells, so a sufficient number of protoplasts were easily obtained from *Aspergillus niger* cells cultured under various conditions. Yeast spheroplasts from *Saccharomyces cerevisiae and Schizosaccharomyces pombe* have also been used in ATAC-seq assays, as previously described (Schep et al. 2015). Furthermore, protoplast preparation could occur with *A. niger* cells cultured under various conditions to decrease experimental variation. In our *Aspergillus niger* ATAC-seq library, we detected the same periodicity pattern in the insert size distribution of sequenced fragments from chromatin as what was observed in a human lymphoblastoid cell line (Buenrostro et al. 2013) and in *Arabidopsis thaliana* (Lu et al. 2017). The *A. niger* ATAC-seq library has an high signal-to-noise ratio, which was controlled by comparing the data to Tn5 integration into naked genomic DNA. The reproducibility of biological and technical replicates in our *A. niger* ATAC-seq protocol was relatively high. In ATAC-seq experiments using mammalian cells and plant cells, typically ∼30-70% of the sequenced reads were aligned to the organellar genomes (Buenrostro et al. 2013; Lu et al. 2017). In our *A. niger* CBS513.88 ATAC-seq results, a significantly higher fraction of reads were mapped to the nuclear genome. These data show that our *Aspergillus* ATAC-seq protocol is an effective method that can be applied to the analysis of other species of filamentous fungi.

We identified accessible chromatin regions from our ATAC-seq data across *Aspergillus* species and compared accessible chromatin regions in the genomes of *A. niger* CBS513.88 and SH-2 strains. Accessible regions tend to overlap much more in comparing regions with the *A. niger* CBS513.88 and SH-2 strains relative to strain-specific regions. These results indicate that many *cis-*elements are common in both strains. The chromatin accessibility regions of *A. niger* CBS513.88 and SH2 were similarly distributed across the various elements of the genome; the chromatin accessibility regions were found in a high proportion at TSSs (∼70%) and in a low proportion at intergenic regions (∼5%). The proportional distribution of accessible regions is different from what is observed in humans (Ackermann et al. 2016), Arabidopsis (Tannenbaum et al. 2018) and tomato (Qiu et al. 2016), which show a high distribution in intergenic regions and gene bodies. This difference may reflect the smaller size of the *A. niger* genome (35 Mb) compared with the larger and less compact genomes of humans (750 Mb), Arabidopsis (125 Mb) and tomato (900 Mb). In addition, TTS also contains a small number of accessibility regions (∼13%), which is similar to what is seen in the mouse (Wu et al. 2016). Therefore, we reasoned that TTSs may also play an important role in the transcriptional regulation of genes and downstream translational control in *A. niger*.

Prior works found that the accessible chromatin regions were positively correlated with gene expression changes of the nearest target genes (Wu et al. 2016; Tannenbaum et al. 2018; Toenhake et al. 2018). We discovered that changing both the culture conditions and deleting TFs can slightly induce differential chromatin accessibility. We also found evidence that a high degree of chromatin changes correlate with transcriptional changes of the nearest target genes through analyses of differential open chromatin (OC) region and the DEGs identified by RNA-seq (Fig. 3B). Interestingly, not all putative target genes with increased open chromatin peaks were upregulated, but a small subset was downregulated (Fig. 3B); these results suggest that regulatory regions could be activated by TF binding and nucleosome depletion, but the result of such regions being accessible could also be gene suppression (Toenhake et al. 2018). Comparative genomics focuses on the comparison of DNA coding regions between different species, revealing the eukaryotic genomic evolution of different *Aspergillus* species (Galagan et al. 2005). In comparing open chromatin regions (non-coding sequences) between *A. niger* and *A. oryzae* species, we discovered that syntenic orthologues of 1564 accessible regions across *A. niger* and *A. oryzae* drive the expression of a collection of target genes that are significantly enriched in housekeeping genes functioning in ribosomes, translation, and energy. The synteny of open chromatin regions between different species may be due to the conservation of non-coding regions (Galagan et al. 2005), indicating that *Aspergillus* species, which act as protein expression cell factories, exhibit the same potential functional elements in protein synthesis regulation.

The accessible chromatin regions identified by ATAC-seq are particularly valuable for the identification of *cis-*regulatory DNA elements (Ackermann et al. 2016; Lu et al. 2017; Tannenbaum et al. 2018; Toenhake et al. 2018), which are bound by regulatory proteins such as transcription factors. We systematically identified genome-wide DNA footprints from candidate accessible regions and performed *de novo* motif enrichment analysis on the footprints from *A. niger* CBS513.88 and SH2 (Fig. 4C). Clustering of TFs by family and analysis of P-values, indicated that the *A. niger* CBS513.88 and SH2 ATAC-seq footprints were enrichment for *de novo* bZIP (motif GCTGAGTCAGCV) and basic helix-loop-helix (motif TCACGTGATC) families (Fig. 4C). The 775 genes predicted as DNA-binding TFs are primarily distributed among 37 classes of regulatory proteins, including 17 bZip family TFs and 10 bHLH family TFs in the *A. niger* genome (Pel et al. 2007; Todd et al. 2014). The bZip family of TFs that have been functionally analysed in *Aspergillus niger* mainly include *hacA* (UPR) (Carvalho et al. 2012) and *cpcA* (amino acid starvation) (Wanke et al. 1997b). The function of other bZip TFs are associated with amino acid starvation, *jlbA* (Strittmatter et al. 2001); oxidative stress, *napA* (Mendoza-Martinez et al. 2017), *atfA* (Sakamoto et al. 2009; Balazs et al. 2010), and *atfB* (Sakamoto et al. 2008); and sulfur metabolism, *cys*-3/*metR* (Amich et al. 2013), and they have mainly been characterized in homologous species, such as *A. nidulans* (Strittmatter et al. 2001; Balazs et al. 2010; Mendoza-Martinez et al. 2017), *A. fumigatus* (Amich et al. 2013) and *A. oryzae* (Sakamoto et al. 2008; Sakamoto et al. 2009). The gene regulatory network of the bZIP family of *Neurospora crassa* (Tian et al. 2011) reveals that the bZIP family of TFs exerts consistent regulatory effects in response to environmental stresses, such as oxidative stress, amino acid starvation, and heavy metal pressure. Functional enrichment of the most proximal gene targeted by the bZip motif, GCTGAGTCAGCV, further demonstrates that this type of TF is associated with transcriptional regulation, which is the first step in regulating the stress response of filamentous fungi to the external environment. The 10 TFs in the bHLH family, including homologous genes of *sclR*/*srbB*, *anbH1*, *srbA*, *palcA*, *INO4,* and *devR*, have not been well characterized in *A. niger*. However, in other species, the bHLH TFs have been postulated to be involved in hyphae growth and differentiation as well as specific metabolic pathways (Caruso et al. 2002; Jin et al. 2011). The GO category of our E-box targeted genes found that they are involved in the biological process of translation, which is identical to the function of the bHLH family proteins that was predicted following *A. nidulans* genome sequence analysis (Galagan et al. 2005). Thus, we reasoned that bHLH TFs in *A. niger* have more unknown functions, and further characterization of bHLH TFs is urgently needed. The bHLH TFs recognize a consensus core element named the E-box (5’-CAVVTG-3’), and the G-box (5’-CACGTG-3’) is the most common form (Jin et al. 2011); further, specific recognition of the sequence relies on flanking sequences (Fisher and Goding 1992; Zhou and O’Shea 2011). The E-box flanking sequences predicted herein contain not only the 5’ T residue but also the 3’ ATC residues. The 5’ T residue flanking the G-box determines major specificity by preventing the binding of other bHLH TFs, such as yeast PHO4 (Zhou and O’Shea 2011), whereas the significance of the 3’ residues is not fully known. The new motif CTGCCTGAGGCA is also a highly enriched motif according to the P-value compared to all *de novo* previously unidentified TF binding motifs. GO functional analysis showed that this new motif CTGCCTGAGGCA drives the expression of target genes, and the GO molecular function categories include the cellular components of “ribosome” and “mitochondrial inner membrane”; the biological processes are related to “translation” and “metabolism”. We constructed a minimal synthetic promoter suitable for filamentous fungi that was based on the yeast minimal promoter (Redden and Alper 2015). We used this promoter to validate the footprint library-driven UAS elements with TF binding site motifs, which provided a new method for *in vivo* validation of *Aspergillus niger* ATAC-seq-predicted binding sites containing TFBs in the absence of other omics data (DNase-seq, ChIP-seq). Our *de novo* motifs identified from enriched from ATAC-seq footprints were able to drive the expression of the targeted gene *in vivo*.

As with DNase I, the cleavage of DNA sequences by the Tn5 transposase is thought to produce pseudo-DNA footprints because of its biased activity (Green et al. 2012; Lu et al. 2017). The digestion frequency analysis of known TF motifs (Fig. 5A) (Lu et al. 2017; Karabacak Calviello et al. 2019; Li et al. 2019) demonstrated a sudden drop in read coverage at the TF site of naked genomic DNA, indicating that Tn5 has a bias in the cleavage of the genome that resulted in pseudo-DNA footprints. Given the enzyme digestion bias, several papers proposed correction methods of TF footprinting (Gusmao et al. 2016; Karabacak Calviello et al. 2019; Li et al. 2019). However, there is no consensus on the significance of correcting DNase I or Tn5 transposase sequence bias due to the different methods used to process Tn5 signals and TF footprinting. In our study, we first applied the TF-defective strain to determine the distribution of Tn5 integration sites around the corresponding TF binding sites to control for Tn5 transposase sequence bias *in vivo.* Tn5 integration patterns around TF binding sites in the absence of the corresponding TF conditions are to the patterns observed in the genomic DNA (Fig. 5A). Prior literature suggests that concave-shaped appearance at the TF motif is known to have transcription factor binding, whereas in *Arabidopsis* footprint prediction, Tn5 digestion of naked DNA produced the same pattern of footprints, which is judged as a pseudo footprint (Lu et al. 2017). The Tn5 digestion frequency of the known motifs in a TF-deficient strain showed that, consistent with the predicted results of the naked DNA, even a TF was knocked out, the same patterns persisted. Therefore, we conclude that the positive and negative DNA strands on both sides of the motif can result in a concave shape that makes it difficult to determine the presence of transcription factor, which can easily produce erroneous results. HINT-ATAC developed by Li et al. (Li et al. 2019) was used to measure the amount of the strand cleavage bias for distinct fragment sizes (All, Nfr, 1N, and +2N) around distinct intervals near the TF binding site, and the results showed intricate strand-specific cleavage patterns relative to TF binding. Strand-specific cleavage patterns around the transcription factor binding site have been reported in multiple species but have been ignored. The strand-specific Tn5 integration patterns around transcription factor binding sites are only observed in our ATAC-seq data from the *A. niger* SH2 wild-type strain, which are caused by TF/DNA interactions and the signal indicating an active TF binding site. By comparing the Tn5 digestion patterns of wild-type strains, TF-deficient strains, and naked DNA footprints, we reasoned that the strand-specific unbalanced pattern of Tn5 digestion on both sides of the footprints can be considered a true DNA footprint.

We further determined the enrichment of known TF binding motifs from ATAC-seq footprints by comparing ATAC-seq data from the *Aspergillus niger* wild-type strain and its TF-defective strains. Footprints for these known TF binding motifs identified by ATAC-seq data were also verified as functional TF binding site *in vivo*. Compared with *A. niger* CBS513.88, the enrichment P-value of the TF motif in *A. niger* SH2 was significantly higher; the most differential values were observed for the CCAAT box, which represents a gene transcriptional activation domain (Wang et al. 2015), and the translation-related bHLH family (Galagan et al. 2005). This suggests that *A. niger* SH2 is a more suitable host for exogenous protein expression (Yin et al. 2014) at the TF level. The TF motif enrichment observed after growth in different carbon source conditions and in comparison to TF-defective strains offer valuable information about TF dynamic responses in addition to the external environment and/or intrinsic regulatory factors. Transcriptome analysis of the known TFs *creA*, *amyR*, *pacC* and *prtT* revealed that these transcription factors are involved in decentralized functions and do not cluster in categories directly related to TF function; this is even true for the pathway-specific transcription factor PrtT, which has a variety of functions that are not directly related to TF function that are clustered. Such results are ubiquitous in other studies using RNA-seq to study the function of transcription factors (Carvalho et al. 2012; Chen et al. 2014; Chung et al. 2014; Zhou et al. 2016), which makes it particularly difficult to confirm the specific regulatory functions of transcription factors; as a result, the targeted GO term of a TF can only be determined by combining phenotypic changes (Chen et al. 2014; Zhou et al. 2016) or ChIP-seq analysis (Chung et al. 2014; de Castro et al. 2014). These functional clustering results are likely to be the effects of cascades of amplified changes caused by TF defects but are not the direct result of TF/DNA interaction. Through the co-regulation analysis of ATAC-seq and RNA-seq of SH2 TF-defective strains (Fig. 6A), we effectively identified gene sets related to the corresponding TFs, and using GO clustering analysis, we screened genes and revealed that the obtained clustering results can accurately reflect the function of the experimentally verified TF (Drysdale et al. 1993; MacCabe et al. 1996; Punt et al. 2008; vanKuyk et al. 2012) (Fig. 6A). Interestingly, we still predicted a different number of corresponding TF binding motifs in the TF knockout strains (delamyR, delprtT, delcreA, delpacC). This result may be caused by three factors: (i) None of these TFs are pioneers that change chromatin accessibility, and the vacancies caused by TF defects are filled by other non-functional proteins. (ii) TF regulation is a complex dynamic process *in vivo* (Qiu et al. 2016; Wu et al. 2016), so that even if a TF was knocked out, there may be different TFs with alternate binding at the same DNA site. For example, the AmyR recognizing motif 5’-CGGN8(C/A)GG-3’ and the PrtT recognizing motif CCG(H)CGG compete for the binding sites in the same target gene during nitrogen and carbon source metabolism regulation (Chen et al. 2014). (iii) These binding site identifications may be the result of a false positive. Unexpectedly, we discovered that footprints contained transcriptional regulators CreA and AreA, and these two factors act not only as repressors but also as activators, which was confirmed by footprint-driven reporter assays *in vivo* (Fig. 6D and E). Our results contradict the fact that it is known that CreA and AreA are repressors for CCR and NCR. It can only be speculated that CreA and AreA may play an unknown activation role in gene transcriptional regulation or that the predicted binding sites contain an activated binding motif for other unknown transcription factors.

These data representing the ATAC-seq landscape of TF footprints will help in the exploration of genome-wide regulation of genes active in filamentous fungi, and the data will help to discover new regulatory factors and their potential sites of action. We demonstrated the usefulness of this resource by testing 25 non-coding DNA sites with predicted TF binding motifs driving reporter gene expression in the host strain: *A. niger* HL-1. Future experiments will be required to validate the functional activity of more noncoding sites and investigate the role of specific TF footprints in controlling gene expression. The sites with *cis*-acting elements provide a large number of basic modules for artificially synthesized promoters, and our ATAC-seq data set will provide a rich resource for the identification of functional TFs in *A. niger*. In the future, ATAC-seq may be effectively applied to study other filamentous fungi whose genomes and transcriptomes are available but whose regulatory mechanisms have not yet been explored.

## Methods

### Strains and media

The specific genotypes of strains used in this study are shown in Supplemental Table S1. *E. coli* Mach 1T1 (Invitrogen, America) was used for molecular cloning and was cultured in Luria-Bertani (LB) medium (1% NaCl, 1% tryptone, and 0.5% yeast extract) with 100 μg/mL ampicillin at 37°C (both solid and liquid) while shaking at 200 rpm. Conidial *Aspergillus niger* CBS513.88 and *Aspergillus oryzae* niaD300 were cultured in PDA medium for spore germination, and they were cultured in DPY medium (2% glucose, 1% peptone, 0.5% yeast extract, 0.5% KH_2_PO_4_ and 0.05% MgSO_4_·7H_2_O) for protoplast preparation. Aconidial *Aspergillus niger* SH2 (Δ*ku*Δ*pyrG*) (Yin et al. 2014) was used as the host for the knock-out of transcription factors and was cultured in modified liquid Czapek-Dox medium (CD) containing the following substances to support mycelial growth: 2% glucose, 0.3% NaNO_3_, 0.1% KH_2_PO_4_, 0.05% MgSO_4_·7H_2_O, 0.2% KCl and 0.01% FeSO_4_·7H_2_O. When the OD reached 2.0, 1 ml of mycelia was inoculated into 100 ml of DPY medium and grown in a thermostatic shaker at 250 rpm and 30°C for 40 h. Mycelia were harvested for protoplast preparation. When necessary, the glucose in DPY was replaced with 2% maltose to prepare MPY medium. *A. niger* HL-1 (Dong et al. 2019) was used as the expression host to produce the enzyme product of the glucose oxidase gene *goxC* (An01g14740) (Guo et al. 2010). Solid CD (2% agarose) with 1% O-dianisidine (dissolved in methanol), 18% glucose, and 90 U/mL horseradish peroxidase was used to assess the colour reaction following *goxC* expression.

### Construction of transcription factor knockout and overexpression strains in *Aspergillus*

Different TFs, *creA* (Drysdale et al. 1993), *amyR* (vanKuyk et al. 2012), *cpcA* (Wanke et al. 1997a), *pacC* (MacCabe et al. 1996), *prtT* (Punt et al. 2008) and *laeA* (Bok and Keller 2004; Bok et al. 2006), were searched on the *Aspergillus* genome database (www.aspgd.com), and homologous arms for gene knockout were designed based on the 1000 bp sequence flanking the CDS region of each gene, which were then amplified with primer sets listed in Supplemental Table S2. The uridine auxotrophic marker *pyrG* from *Aspergillus nidulans* was used as a screening marker for TF knockout. The upstream and downstream homologous arms of each TF obtained by PCR amplification were introduced along with the *pyrG* marker into a pMD20 T-Vector (TaKaRa, Japan) by NEBuilder (New England Biolabs, America), to obtain TF knockout vectors (pdelcreA, pdelamyR, pdelcpcA, pdelpacC, pdelprtT and pdellaeA). The knockout cassettes that were linearized with *XbaI*, *EcoRI*, *ApaI*, *EcoRI*, *EcoRI* and *BamHI* were introduced into *A. niger* SH2 and *A. oryzae* niaD300 by PEG-mediated protoplast transformation. The cassettes for *laeA* overexpression digested with *ApaI* were transformed into *A. oryzae* niaD300 to yield the OElaeA strain. The TF knockout mutants in which homologous integration occurred were analysed by PCR amplification for upstream and downstream localization, and they were further validated by qRT-PCR (Supplemental Fig. S1). The primers used in this study are listed in Supplemental Table S2.

### ATAC-seq library preparation

The *A. niger* strain CBS513.88, SH2 (including TF deletion strains) and *A. oryzae* niaD300 (including *laeA* deletion and overexpression mutants) were grown to mid-log phase in DPY medium or MPY medium. Then, 1 g of mycelia per library were harvested using miracloth, and they were then rinsed with cold water and 0.8 M NaCl. Protoplasts were prepared by incubating mycelial samples in enzymatic lysis buffer composed of 0.8 M NaCl, 2% cellulase, 1% helicase, 1% lyticase and 0.5% lysozyme, which were dissolved in the corresponding medium and were then filtered with miracloth and washed with cold STC buffer (10 mM Tri-HCl pH=7.5, 1.2 M sorbitol, 50 mM CaCl_2_). Subsequently, 5×10^4^ protoplasts (cells) were incubated with 200 μl of lysis buffer (10 mM Tris-HCl, pH 7.4, 10 mM NaCl, 3 mM MgCl_2_, 0.05% (v/v) IGEPAL CA-630, 1 mM PMSF and 1xPIC (EDTA free)) for 10 min at 4°C, washed with lysis buffer (without IGEPAL CA-630), and then incubated with 5 μl of Tn5 Transposase of TTE Mix V50 in 45 μL of 1 x TTBL buffer (TruePrep® DNA Library Prep kit V2 for Illumina®, Vazyme, China) at 37°C for 30 min. PCR was subsequently performed as previously described (Buenrostro et al. 2013). Libraries were purified with AMPure beads (Agencourt) to remove contaminating primer dimers. All libraries were sequenced with 50 bp paired-end reads on an Illumina Hiseq 2500 platform. For the control (*A. niger* CBS513.88 naked genome DNA), 5 ng of genomic DNA was used for the Tn5 transposase reaction.

### RNA extraction, qRT-PCR, and RNA-seq analysis

A small portion of the mycelium used to prepare the ATAC-seq library was left for RNA extraction. Total RNA was isolated using a HiPure Fungal RNA Kit (Magen, China). Reverse transcription was performed with a PrimeScript RT-PCR Kit (TaKaRa, Japan). Quantitative real-time PCR (qRT-PCR) was performed using an ABI 7500 Fast Real-Time PCR System to analyse TFs and the report gene *goxC*; the primers used are listed in Supplemental Table S2. The resulting libraries were sequenced at the Beijing Genomics Institute (BGI) with 50 bp single-end reads on the BGISEQ-500. RNA quality was assessed with an Agilent Bioanalyzer 2100 system to confirm that all samples had an RNA integrity value (above 7.0). All samples were prepared in duplicate. To obtain FASTQ format clean reads, all generated raw sequencing reads were filtered to remove reads with adaptors, reads in which unknown bases were greater than 10%, and low-quality reads. After filtering, clean reads were mapped to the genome and cDNA using HISAT (Kim et al. 2015) and Bowtie2 (Langmead and Salzberg 2012). The gene expression level was quantified by RSEM (Li and Dewey 2011). The DESeq2 method (Love et al. 2014) was used to screen differentially expressed genes between the two groups.

### Mapping and normalization of ATAC-seq data

After removing adaptors using Trimmomatic (Bolger et al. 2014), 50 bp paired-end ATAC-seq reads were mapped to the *Aspergillus* genome (Ensembl/ASM285 for *A. niger* and Ensemble/ASM18445 for *A. oryzae*) using Bowtie2 (Langmead and Salzberg 2012) with parameters “--trim5 5 --trim3 15” and other default parameters. All reads aligned to the + strand and the – strand were offset by ±5 bp, respectively, due to the Tn5 transposase introducing two cuts that were separated by 9 bp (Adey et al. 2010). Mapped reads of SAM output were converted to a BAM format and sorted by Samtools (Li et al. 2009). Duplicate reads were removed using the default parameters of the Picard tools MarkDuplicates program (http://broadinstitute.github.io/picard/).

To visualize mapped reads, BAM format files were converted to bigwig format using the *bamCoverage* tool in deepTools2 (Ramirez et al. 2016) with a bin size of 1 bp and with RPKM normalization. Heatmaps and average plots from the ATAC-seq data matrix were also generated using the ‘*computeMatrix*’, ‘*plotHeatmap*’ and ‘*plotProfile*’ functions in the deepTools2 package. Genome browser images were made using the integrative Genomics Viewer (IGV) (Thorvaldsdottir et al. 2013) and bigwig files that were processed as described above. Fragments of the desired length were extracted from SAM files.

### Peak calling identified accessible chromatin regions

Accessible regions and peaks of each sample were identified using MACS2 (Zhang et al. 2008) with the following parameters: -f BAMPE --nomodel --shift 100 --extsize 200. The narrowPeak files from the MACS2 output were used for further analysis. The *annotatePeaks.pl* tool of the HOMER program (Heinz et al. 2010) was used with the default parameters to annotate the location of the identified peaks overlapping with genomic features, the TSS (transcription start site), the TTS (transcription termination site), exons, introns, and intergenic regions, all of which were annotated in the Ensemble *Aspergillus niger* CBS513.88 genome. Peaks that were 1 kb upstream of the TSS were associated with the nearest genes. These genes were then analysed for overrepresented gene ontology and KEGG pathway using FungiFun 2.2.8 BETA (Priebe et al. 2015) (https://sbi.hki-jena.de/fungifun/).

### Analysis of differential chromatin accessibility

To identify differentially mapped reads, we used Diffbind (Ross-Innes et al. 2012). We used the processed ATAC-seq alignment bam file from Bowtie2 and the narrowPeak file from MACS2 for each sample. MA plots (log2-fold change vs. mean average) were used to visualize changes in chromatin accessibility for all peaks.

### Identification of footprints

Footprints were identified with pyDNase (Piper et al. 2013) using the default parameters. The Wellington footprinting algorithms (Piper et al. 2013) in pyDNase searched for TF footprints in areas of the genome that MACS2 identified as peaks. Wellington applied a beta-binomial distribution to estimate footprints and elevated the strand-specific activity of DNase or Tn5 transposase around transcription factor footprints among accessible chromatin regions.

### Design and synthesis of a minimal *Aspergillus* promoter for functional verification of CREs *in vivo*

With reference to the yeast minimal promoter screening system (Redden and Alper 2015), we designed a minimal *Aspergillus* promoter for functional verification of CREs *in vivo* (Supplemental Fig. S2A). The 67 bp 5’UTR of the *Aspergillus nidulans TPI* gene was used as a TSS (McKnight et al. 1986). The TATA box and the TSS are separated by a 29 bp distance from the TSS in the *amyB* (An05g02100) promoter of *A. niger*, which facilitates successful loading of the transcription initiation complex, thereby initiating transcription by RNA polymerase. To ensure that the TATA box can be successfully opened during the transcription process, we manually added 8 AT-enriched sequences in front of the TA box (Redden and Alper 2015) to reduce the bond energy of the DNA. These elements were linked together by fusion PCR amplification to form a core promoter region (Supplemental Fig. S2A). The glucose oxidase *goxC* (An01g14740) was used as the reporter gene, which generates coloured colonies (Supplemental Fig. S2B) (Guo et al. 2010). To further refine the entire reporting system, the selection marker *pyrG* was integrated before the footprints to separate the footprints from other genomic sequences, ensuring that transcription of the core sequence was not affected by other sequences. The footprints contained a transcription factor binding site (TFBS) that helped stabilize RNAP to enhance transcription efficiency. By constantly changing footprints and monitoring the transcription of downstream reporter genes, it was possible to identify whether the footprints containing the TFBs function *in vivo*. The 1 kb flanking sequence of the glucoamylase gene (*glaA*) acts as a homologous arm, and the aim was that the sequence successfully integrated into the glucoamylase gene locus, ensuring that all open reading frames were at the same position in the genome (Rantasalo et al. 2018).

### Discovery of *de novo* transcription factor motifs

ATAC-seq footprints that were identified from each sample were used to discover *de novo* transcription factor motifs. The boundaries of the start and stop coordinates of ATAC-seq footprints were lengthened by 10 bp upstream and downstream. We then used the *findMotifsGenome.pl* tool of the HOMER program to identify over-represented transcription factor sites (transcription factor motifs). Using the *annotatePeaks.pl* tool of HOMER with the –m parameter, HOMER *de novo* motifs were then used to search the original set of sequences to identify all instances of motifs.

### Analysis of known motif enrichment within footprints

The known motif files in the HOMER format were created manually with the *seq2profile.pl* tool of HOMER. The known motifs were then used to search the footprints of each sample to identify all instances of motifs using the *annotatePeaks.pl* tool of HOMER with the –m parameter.

## Data access

The raw ATAC-seq sequencing data have been deposited in the NCBI SRA database under the accession number PRJNA566304 and PRJNA587805. The raw RNA-seq sequencing data have been deposited in the NCBI SRA database under the accession number PRJNA588127.

## Acknowledgements

This work was supported by the National Natural Science Foundation of China [grant number 31871736, 31870024].

## Disclosure declaration

None conflicts of interest.

